# Competition effects regulating the composition of the microRNA pool

**DOI:** 10.1101/2024.05.28.596158

**Authors:** Sofia B. Raak, Jonathan G. Hanley, Cian O’Donnell

## Abstract

MicroRNAS (miRNAs) are short non-coding RNAs that can repress mRNA translation to regulate protein synthesis. During their maturation, multiple types of pre-miRNAs compete for a shared pool of the enzyme Dicer. It is unknown how this competition for a shared resource influences the relative expression of mature miRNAs. We study this process in a computational model of pre-miRNA maturation, fitted to *in vitro* Drosophila S2 cell data. We find that those pre-miRNAs which efficiently interact with Dicer outcompete other pre-miRNAs, when Dicer is scarce. To test our model predictions, we re-analysed previously published *ex vivo* mouse striatum data with reduced *Dicer1* expression. We calculated a proxy measure for pre-miRNA affinity to TRBP (a protein which loads pre-miRNAs to Dicer). This measure well-predicted mature miRNA levels in the data, validating our assumptions. We used this as a basis to test the the model’s predictions through further analysis of the data. We found that pre-miRNAs with strong TRBP association are over-represented in competition conditions, consistent with the modelling. Finally using further simulations, we discovered that pre-miRNAs with low maturation rates can affect the mature miRNA pool via competition among pre-miRNAs. Overall, this work presents evidence of pre-miRNA competition regulating the composition of mature miRNAs.

## Introduction

MicroRNAs (miRNAs) are small non-coding RNAs that inhibit protein translation via the RNA-induced gene silencing complex (RISC). MiRNAs are synthesised in the nucleus by RNA polymerase II/III as primary-miRNAs (pri-miRNAs), which are then cleaved by Drosha/DGCR8 to form precursor miRNas (Results; Figure 1 A). Pre-miRNAs are exported into the cytosol via Exportin-5 and transported to sites of local inhibition of protein translation, such as neuronal dendrites, where they are loaded onto Dicer, which cleaves the characteristic hairpin-loop structure to produce mature miRNA. The double-stranded miRNA is then loaded onto Argonaut proteins (Ago), which finish the maturation by ejecting the passenger strand to leave a single-stranded miRNA bound to Ago. The Ago-bound miRNA can subsequently undergo complementary base-pairing with target mRNAs and trigger RISC assembly, leading to silencing of protein translation.

**Fig. 1:**
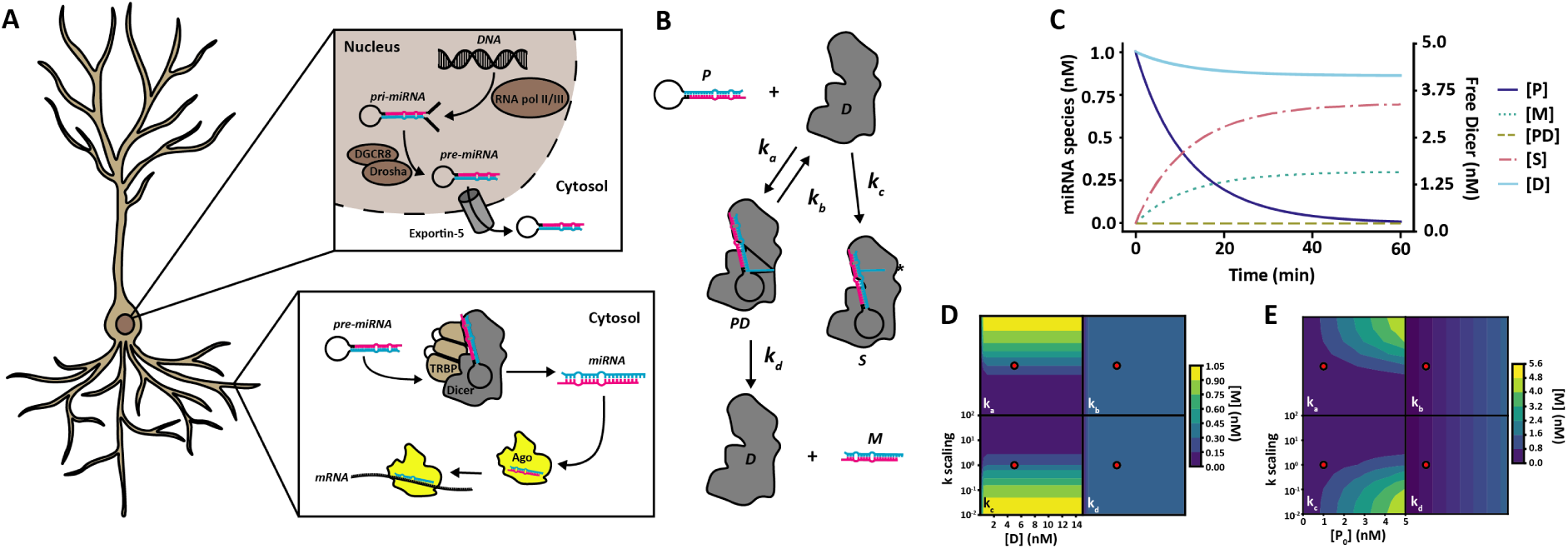
MicroRNA maturation in the neuron and model design. **A** MiRNA maturation and function in the neuron. MiRNA is synthesised in the nucleus as pri-miRNA by RNApol II/III as a long single-stranded RNA molecule with a central hairpin loop. Pri-miRNA is then cleaved by the Drosha/DGCR8 complex to release the hairpin loop as pre-miRNA. Pre-miRNA is then transported out of the nucleus into the cytosol by the Exportin-5 complex. In the cytosol, pre-miRNA can be transported into dendrites where Dicer, assisted by e.g. TRBP, binds pre-miRNA. Dicer cleaves the pre-miRNA by the loop structure and double-stranded mature miRNA can then be loaded into Argonaute proteins, where the passenger strand is ejected and the remaining single-stranded miRNA can form complimentary base-pair binding with target mRNA for targeted repression of local protein translation. **B** Model diagram of pre-miRNA maturation. In the computational model, pre-miRNA (*P*) can associate with Dicer (*D*) to form a transient Dicer-pre-miRNA complex (*PD*) with rate *k_a_*. The Dicer-pre-miRNA complex can either deteriorate back to free pre-miRNA and Dicer at rate *k_b_*, or go through pre-miRNA maturation at rate *k_d_* to form free Dicer and mature miRNA (*M*). Alternatively, free pre-miRNA and Dicer can associate to form a stalled complex of Dicer-pre-miRNA (*S*) at rate *k_c_*, which permanently binds pre-miRNA and Dicer in the system. **C** Dynamics of species concentration in the model. As time increases, the concentration of free Dicer ([D]) and pre-miRNA ([P]) reduces while the concentration of mature miRNA ([M]) and stalled pre-miRNA and Dicer ([S]) increases. The transient pre-miRNA Dicer complex [PD] is highly unstable and does not accumulate in the system. Parameter values for this simulation are given in Table 1. **D** Effects of varying Dicer concentration and reaction rates on miRNA concentration at steady state. Reaction rates were scaled from 10^-2^ to 10^2^ times the value obtained from data fitting (recorded in Table 1; wild-type parameters used, indicated by red dot) while Dicer concentration was varied from 0.01 nM to 15 nM. Increasing reaction rate for association (k_a_) leads to an increase in mature miRNA concentration at steady state, whereas increasing stalling (k_c_) leads to a decrease. No effect is seen when changing dissociation (k_b_) or dicing (kd) rates. The effects of varying Dicer concentrations in the system are only notable at very low Dicer concentrations, where mature miRNA concentration at steady state reduces, confirming that there is an abundance of Dicer available in the system used by [20]. **E** Effects of varying reaction rates and initial pre-miRNA concentration ([P_0_]) at steady state. Reaction rates were scaled as for D and initial pre-miRNA concentration varied between 0 and 5 nM. As in D, increasing k_a_ increases the amount of mature miRNA, whereas increasing k_c_ leads to a decrease at steady state. Increasing the pre-miRNA concentration modestly increases mature miRNA regardless of variation in dissociation (k_b_) or dicing (k_d_) rates. Red dots represent default values obtained from optimisation and used in C.

MiRNAs are of particular importance in regulating gene expression in dendrites due to the size and morphology of neurons. For example, in a cortical pyramidal neuron, the soma is typically around 20µm in length, whereas some dendrites can extend hundreds of micrometres [1] and when considering the entire dendritic arborisation, the total length of dendrite for a single neuron can reach tens of millimetres [1, 2]. Given these large distances, it seems likely that neurons must use local mechanisms to control the spatial pattern of protein expression, rather than orchestrating control completely from the nucleus. Control and maintenance of dendritic pools of mRNA transcripts offer an elegant solution to highly localised and highly specific translational control of the post-synaptic proteome via RNA-induced gene silencing by miRNAs [3]. MiRNAs therefore play an important role in synaptic function and plasticity in the brain.

Competition is a recurring theme at all levels of biology, from competition between species and individual organisms to the competition for resources on the molecular level within the cell. Competition effects in biosynthesis has been heavily studied using computational models (reviewed by [4]). Early studies on prokaryotic transcription highlighted key parameters governing competition between sigma factors for RNA polymerases [5, 6]. Mauri and Klumpp’s (2014) model in particular was structurally similar to the model of competitive miRNA maturation we study here. Other studies examined the role of competition in protein translation, for example due to limited availability of ribosomes or tRNAs [7–9]. Several computational modelling studies have also explored the effects of competition between miRNAs and mRNAs [10–13], consistent with *in vitro* experiments [14]. Collectively these insights are of great importance for synthetic biology applications, where expression of exogeneous genes can put strain on endogeneous biosynthesis machinery [15–17]. For example, miRNA-mRNA competition can affect noise in synthetic gene circuits [18]. In tissues with high pre-miRNA expression levels, such as the brain, where up to 70% of known miRNAs have been detected [19], it is reasonable to assume that a large number of pre-miRNAs with different Dicer affinities and maturation efficiencies are competing for a limited amount of available Dicer. If this is the case, pre-miRNA competition for Dicer may indirectly regulate the composition of the mature miRNA pool.

For competition to be meaningful, individual components of a system must display distinct and diverse attributes. Among pre-miRNAs, both sequence and structural characteristics have been linked to the efficiency of maturation. Tsutsumi *et al.* (2011) [20] showed that *Drosophila* Dicer1 is more efficient at cleaving pre-miRNAs with a large loop size *in vitro*. Work by Luo *et al.* in HEK293T cells expressing recombinant Dicer also showed a preference of Dicer towards pre-miRNAs with a large loop structure and strong binding in the stem region [21], suggesting that loop size may also play a role in regulating pre-miRNA association *in vivo*. More recently, Lee *et al.* (2023) [22] identified a conserved sequence motif, the GYM motif, which is recognised by the human Dicer1 double-stranded RNA binding domain and is associated with highly efficient cleavage of specific pre-miRNAs. These are all provide different advantages for select pre-miRNAs in maturation and might drive an over-representation of specific mature miRNAs in conditions with reduced Dicer availability.

As discussed above, miRNA competition has been studied before, but most studies have focused on the competition between mature miRNAs, or miRNAs and non-coding competing endogenous RNAs, for the same mRNA targets (see [23–25] for some examples). To our knowledge, competition between different pre-miRNA species for proteins in the miRNA maturation pathway has not been reported previously. Here, we present a simple computational model of pre-miRNA maturation and pre-miRNA competition for Dicer based on mass-action kinetics (Results; Figure 1B). The model predicts that pre-miRNAs with both a high rate of association with Dicer and efficient dicing rates have a competitive advantage over other pre-miRNAs in systems with both abundant and severely reduced Dicer. Based on our model predictions, we identify pre-miRNA competition for Dicer *in vivo* from previously published experimental data [22]. Our work highlights the non-specific effects of pre-miRNA competition for Dicer on the global miRNA pool.

## Results

### MiRNAs with a fast association rate to Dicer display robust maturation levels in competitive conditions

We designed a minimal model of pre-miRNA maturation (Figure 1B) that could account for the dynamics of mature miRNA production in previously published *in vitro* time series data [20] (See Methods). To summarise the model briefly, a pool of pre-miRNA can reversibly bind with free Dicer, then go through a subsequent maturation step, resulting in the conversion of pre-miRNA to mature miRNA and the release of Dicer back to the free pool. Alternatively, pre-miRNA could irreversibly bind with Dicer forming a stalled complex. We initialised the model with pre-miRNA and Dicer only.

In neurons, pre-miRNA maturation can take place hundreds of µm from the soma where transcription and initial maturation from pri-miRNA to pre-miRNA takes place (Results; Figure 1A). Experimental data has shown that mRNA transcripts can be transported across dendrites in bursts of speeds from 0.5-5 µms^-1^ between short pauses of being stationary [26,27]. Under the assumption that pre-miRNAs are transported at a similar rate in a similar fashion to mRNAs we decided to omit pre-miRNA replenishing from our model.

During a 60 minute simulation of the model (Figure 1C), the concentration of free Dicer and pre-miRNA drops over time while the concentration of pre-miRNA complexed with Dicer increases, along with mature miRNA and stalled pre-miRNA and Dicer. At the end of the simulation, no free pre-miRNA remains and the mature miRNA concentration has reached a plateau. To better understand the system, a model with a single miRNA modelled on the reaction dynamics of wild-type *let-7* [20] was used to vary reaction rates, Dicer concentration (Figure 1D) and initial pre-miRNA concentration (Figure 1E). The model was allowed to run until a steady state was achieved, after which the mature miRNA concentration was extracted and plotted against the reaction rates and Dicer or intial pre-miRNA concentration respectively. As expected, increasing the association rate (k_a_) leads to an increase in final mature miRNA concentration, with the reverse seen for the stalling rates (k_c_). Due to an abundance of Dicer in the system, increasing Dicer concentration does not lead to an increase in miRNA concentration (Figure 1D), as the theoretical maximum miRNA concentration is reached at a low level. Increasing the rate of dissociation (k_b_) or dicing rate (k_d_) did not significantly change the final mature miRNA, suggesting that these reaction steps do not individually significantly alter the reaction dynamics of the system. When varying pre-miRNA concentrations instead of Dicer a similar pattern is seen for each varied reaction rate, with increased mature miRNA with increased reaction rate for species where association rate (k_a_) was varied. Some increase was also seen at low stalling rate (k_b_) and high pre-miRNA concentrations. We also observed modest increases in mature miRNA with increased pre-miRNA for species with varying dissociation and dicing rates (k_b_ and k_d_), highlighting that pre-miRNA availability is more important in determining the final miRNA concentration than dissociation and dicing rate or Dicer availability (Figure 1E).

Next, we used this model in a set of simulation experiments to investigate how multiple different pre-miRNA types might compete for a shared pool of Dicer. There are over 1900 types of mature human miRNAs recorded in miRBase [28], the online repository of identified miRNAs. While miRNAs are highly localised, many different miRNA species may still compete for a shared Dicer pool. The expression levels of these miRNAs vary substantially [29], implying that they have heterogeneous Dicer affinities and maturation rates.

In our model, four distinct reaction rates can affect pre-miRNA maturation: association to Dicer (*k_a_*), dissociation from Dicer without maturing (*k_b_*), irreversible association to Dicer leading to a stalled complex (*k_c_*), and maturation through dicing (*k_d_*). This allowed us to dissect the role of each stage in pre-miRNA maturation by varying each reaction rate independently. To achieve this, we designed 8 theoretical species of pre-miRNA (Figure 2A). We increased each parameter value in turn either 10-fold (high) or 20-fold (2x high) from the optimised values to investigate what characteristics can be expected to confer advantages in competitive environments. As the actual concentration of Dicer in a cellular environment is unknown, we ran a series of simulations with 1nM of each pre-miRNA species present and a range of 0.01 to 8 nM Dicer available until a steady state was reached (Figure 2C-H). In order to reach steady state for each condition and each pre-miRNA, the simulation was run for 3000 minutes and the mature miRNA concentration at the end of the simulation used. Since there was 8 nM total pre-miRNA, we should expect to see competition effects emerge at Dicer concentrations between 0–8 nM. The exact conditions for competition also depends on the particular set of pre-miRNA binding affinities.

**Fig. 2:**
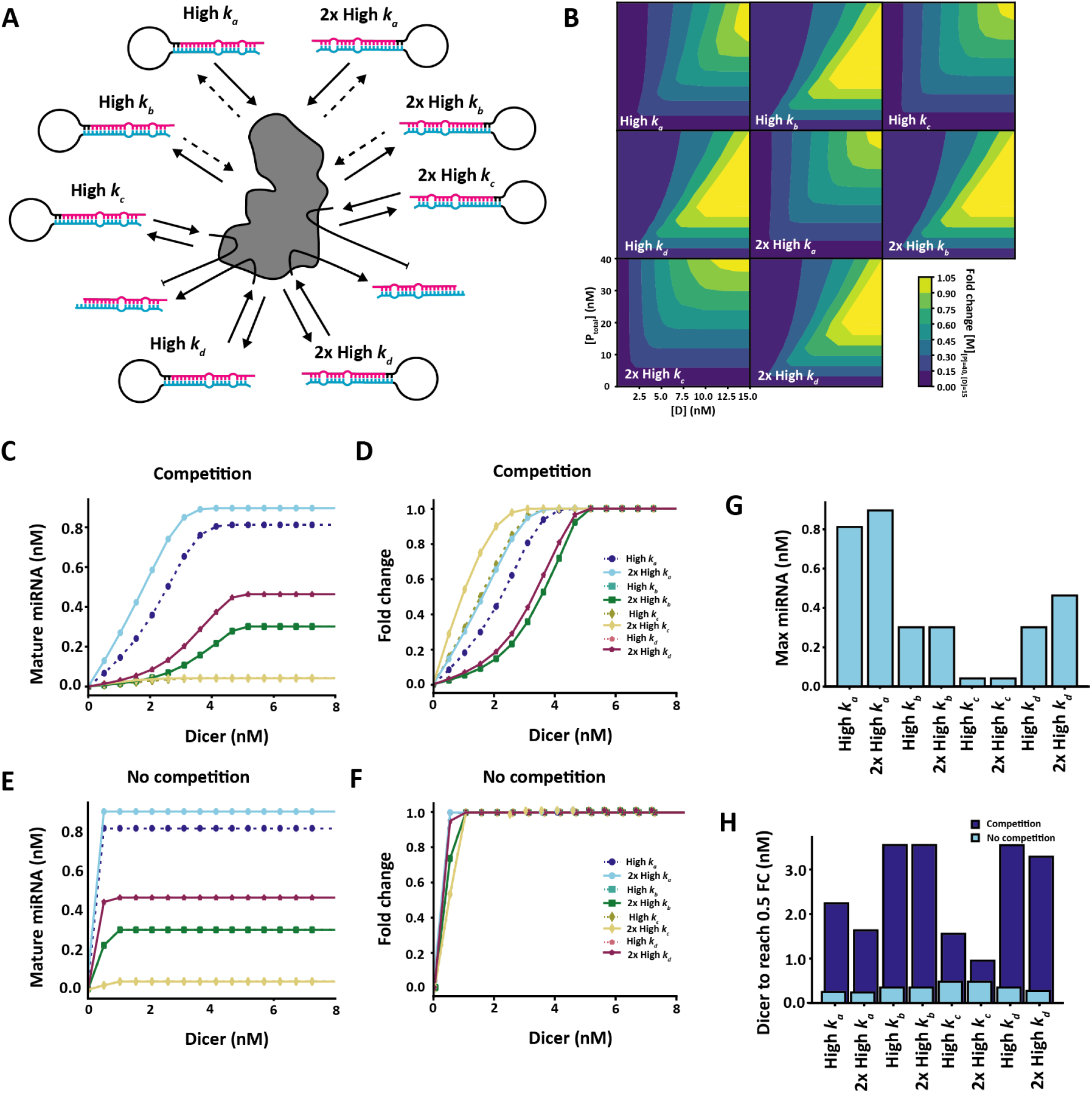
Competition effects between multiple pre-miRNA species. **A** Competition diagram. Each reaction rate was increased 10-fold (high) or 20-fold (2x high) per pre-miRNA species.**B** Effects of varying initial pre-miRNA in the system on competition. The pre-miRNA concentration for each species was varied uniformly along with Dicer concentration. Each simulation was run until steady state was reached and the fold change in mature miRNA for each species calculated on the condition with 5 nM initial pre-miRNA (40 nM total pre-miRNA in the system) and 15 nM Dicer. Increasing Dicer leads to a general increase in fold change for each pre-miRNA, however an increase in pre-miRNA concentration leads to a decrease in fold change among pre-miRNAs with fast dissociation rates (high k_b_ and 2x high k_b_). **C-D** Effects of varying Dicer availability on final mature miRNA concentration in the presence (C) and absence (D) of competition. **E-F** Effects of varying Dicer availability on miRNA maturation fold change, as calculated based on 8n M Dicer availability in the presence (E) and absence (F) of competition. **G** Final mature miRNA concentration at 10 nM Dicer availability. **H** Minimum Dicer needed to reach 0.5 fold-change for each pre-miRNA species in the presence and absence of competition. Final mature miRNA concentration obtained at steady state.

In a competitive environment with a single Dicer pool, high rates of association (*k_a_*) leads to a higher amount of mature miRNA (Figure 2C). This is also true in the absence of competition (Figure 2E), however in competitive regimes, when Dicer concentration is low, slightly higher levels of pre-miRNAs with fast association rates reach the mature state compared to pre-miRNAs with high rates of dicing or maturation (*k_d_*). When investigating fold-change from the simulation with 8 nM Dicer, a high rate of association (*k_a_*) is also highly advantageous both in presence and absence of competition (Figure 2D and F), though not as resistant to competition as pre-miRNAs with a high stalling rate (*k_c_*). In contrast, pre-miRNAs with a high dicing rate (*k_d_*) and high level of stalling (*k_c_*) are almost equally sensitive to a drop in fold change than pre-miRNAs with high association rates (Figure 2D and H), despite a high dicing rate being highly advantageous in pre-miRNA maturation (Figure 2C-F). When Dicer is available in abundance and competition is negligible, a high association rate (*k_a_*) also leads to a higher rate of pre-miRNA maturation than a high dicing rate (*k_d_*, Figure 2G). This suggests that pre-miRNAs with features that promote Dicer association provide both a competitive advantage in environments with reduced Dicer availability and reach the highest level of maturation efficiency. This general effect is preserved over a range of pre-miRNA concentrations in the system. When varying the total initial pre-miRNA in the system by scaling all pre-miRNAs simultaneously between 0 and 5 nM, pre-miRNAs with high association rates (k_a_ remain highly expressed (Figure 2B). In contrast, increasing the amount of initial pre-miRNA in the system leads to a notable decrease in pre-miRNAs with a high dissociation rate (k_b_; Figure 2B), confirming that they are most sensitive to competition effects between miRNAs. This non-monotonic dependence of low-Dicer-affinity miRNAs to global pre-miRNA abundance is an interesting prediction of the model.

### Signatures of pre-miRNA competition for Dicer in experimental data

To test whether any evidence of pre-miRNA competition effects can be detected in experimental data, we next investigated the characteristics of miRNA sequences from the YAC128 mouse model of Huntington’s disease where *Dicer1* mRNA expression levels in the YAC128 mice (expressing transgenic human *HTT* with 100-120 glutamine repeats) have been reported to be reduced by half compared to wild-type mice (which express native mouse *Htt* only) (Figure 4D), while mRNA expression levels of proteins in the pri-miRNA processing machinery or pre-miRNA export were unaffected [30]. We hypothesised that, since Dicer expression was reduced in the Huntington’s model, there should be stronger competition between the pre-miRNAs in that scenario, compared to wild-type animals where Dicer was more abundant. Our strategy was as follows. Our simulation results above predicted that pre-miRNAs with stronger Dicer affinity should out-compete those with low Dicer affinity. Therefore, we aimed to identify some proxy measure that correlates with Dicer affinity, calculate that quantity for each pri-miRNA, and ask if it is predictive of the fold-change in mature miRNA expression in the Huntington model relative to wild-type. The specific prediction was that high-Dicer-affinity miRNAs should show a lower fold-drop in expression than low-Dicer-affinity miRNAs (red line in Figure 3C). In contrast, a lack of competition for Dicer should result in a flat fold-change in miRNA expression, independent of Dicer affinity (dashed blue line in Figure 3C).

**Fig. 3:**
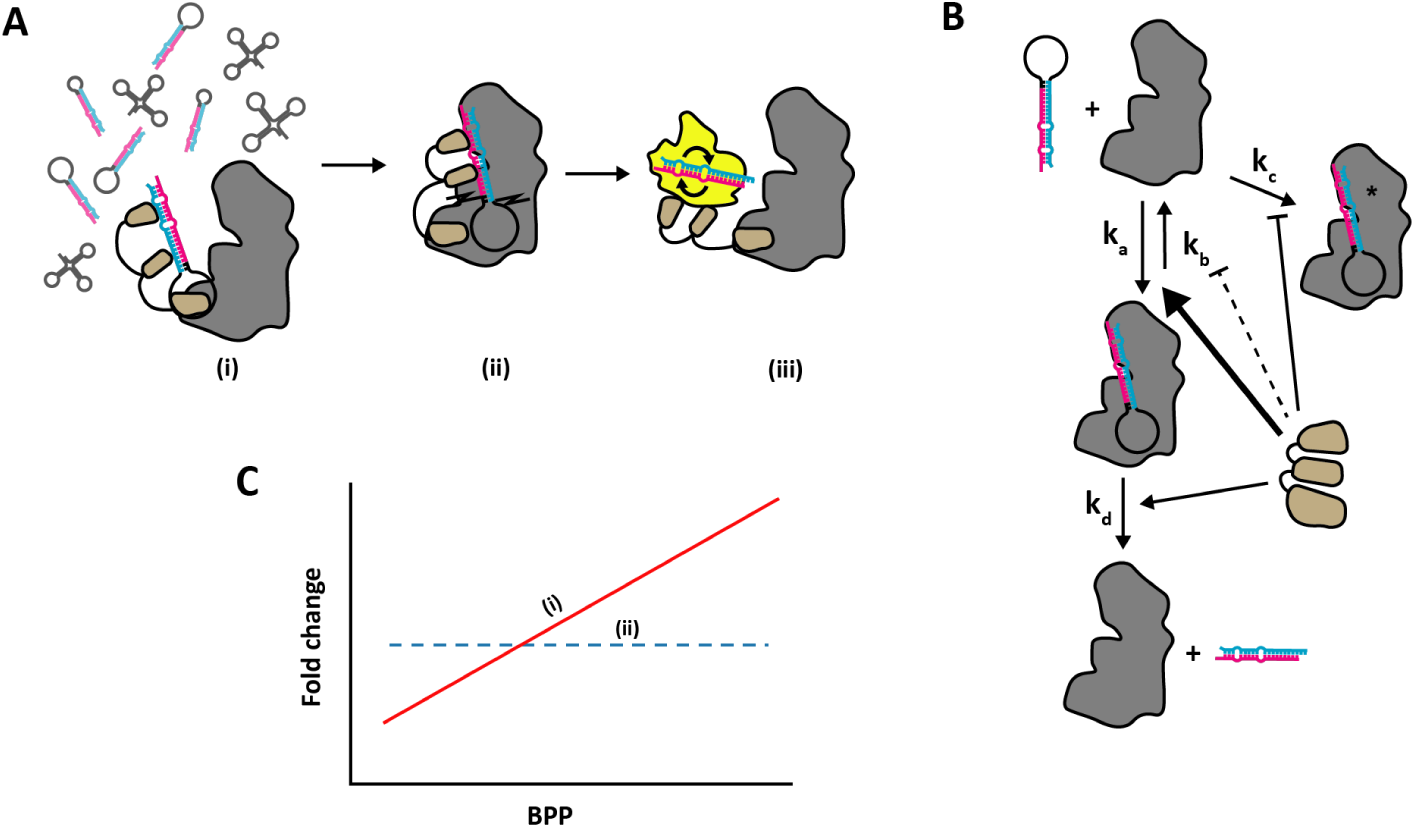
Strategy for identifying competition in experimental data. **A** Role of TRBP in pre-miRNA maturation. TRBP forms a complex with Dicer where it aids in recognition and loading of pre-miRNA onto the catalytic site of Dicer (i) in crowded environments [34]. TRBP also aids in determining the appropriate cleavage site (ii) and influences strand selction by Ago (iii) [32]. **B** Proposed effects of TRBP on model parameters. TRBP association corresponds to boosted association rate *k_a_* and cleavage rate *k_d_*, while the dissociation rate *k_b_* and stalling rate *k_c_* is moderately reduced. **C** Predicted effects in data. If competition effects are present, we expect that a fast association rate (*k_a_*) should have the biggest effect. A strong base-pair binding probability (BPP) in the stem region should be associated with stronger TRBP binding and therefore more efficient Dicer loading. We can calculate base-pair binding probabilities for sequenced miRNAs based on available structures and correlate BPP with miRNA expression levels in the wild-type and reduced *Dicer1* (YAC128) mice respectively. We expect a positive correlation between BPP and miRNA expression if pre-miRNAs with high BPP have a competitive advantage in miRNA maturation (i; red solid line), with a stronger correlation in the YAC128 mice where *Dicer1* expression is reduced. If no competition is present, or fast Dicer association is not providing a significant advantage in miRNA maturation, we expect no relationship between BPP and miRNA expression (ii; blue dotted line).

**Fig. 4:**
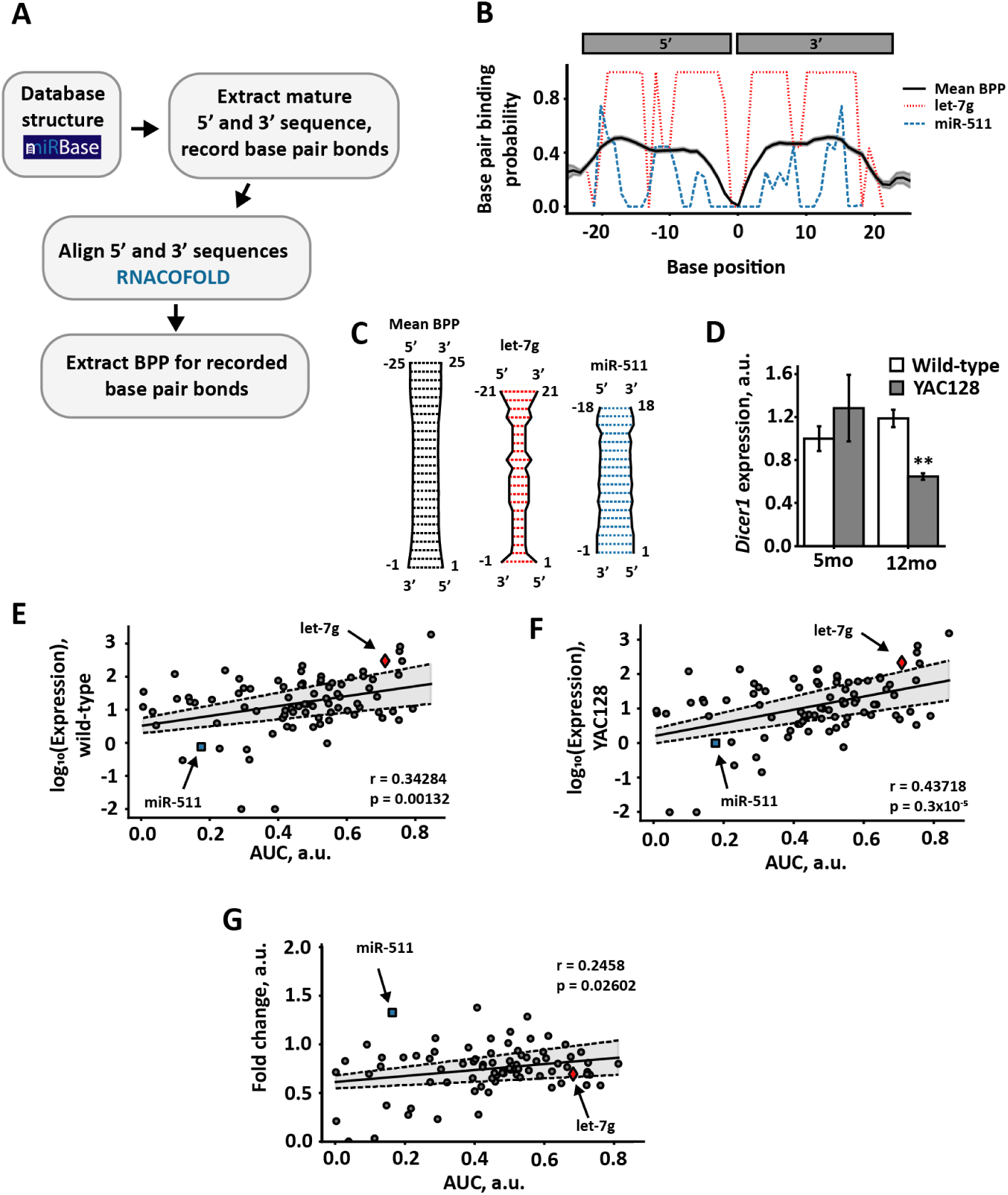
Analysis of pre-miRNA structures suggest maturation advantage of pre-miRNAs with strong associations to Dicer in an HD mouse model. **A** Bioinformatics approach. Pre-miRNA hairpin structures were obtained from mirBase [28] and the mature sequence with recorded base pair bonds extracted. Mature 5’ and 3’ sequences were then aligned using RNACofold v. 2.4.13 from the ViennaRNA 2.0 package [35]. The base pair binding probability (BPP) was then extracted for the recorded base pair bonds. Where no base pair bond was recorded the mean BPP for the i_th_ 5’ nucleotide was used. **B** Subset of base pair binding probabilities. Black solid line represents mean base pair binding probabilities across analysed miRNAs, with shaded area representing standard error of the mean. Red dashed line represents an example miRNA (let-7g) with higher mean BPP, blue dotted line an example miRNA (miR-511) with lower than mean BPP. Position on x-axis denotes nucleotide position, with negative numbers referring to position on the 5’ strand and positive numbers position on the 3’ strand. The centre arbitrarily assigned 0 corresponds to the cleavage site. **C** Base pair binding probabilities of mature miRNA sequences. Graphical representation of subset of the mean base pair binding probabilities of all miRNAs (black) and an example miRNA with higher (let-7g, red) and lower (miR-511, blue) mean BPP. **D** *Dicer1* mRNA expression in wild-type and YAC128 mice at 5 months and 12 months of age as determined by qPCR. P**<0.01 as determined by Mann-Whitney U-test, error bars represent standard deviation. Figure adapted from [30]. **E-F** Pearson correlation of miRNA expression in 12 month old wild-type (**E**) and YAC128 (**F**) mice with BPP area under curve (AUC). **G** Pearson correlation of fold change between 12 month old YAC128 and wild-type miRNA expression with BPP AUC. In **E-G** solid line represents least squares linear regression, with shaded area calculated from the standard error of the intercept and gradient. In **G** outliers above fold change 4 were removed.

We used the miRNA sequencing data from 12-month old YAC128 and wild-type mice produced in Lee *et al.* (2011) [30] to look for pre-miRNA features that might influence maturation. The RNA-loading complex (RLC) protein TRBP has been shown to promote efficient loading and processing of pre-miRNAs in crowded environments [31], as well as promoting cleavage of the hairpin loop at the correct site and strand selection during loading onto Ago proteins [32] (Figure 3A). Additionally Takahashi *et al.* (2018) [33] showed that TRBP preferentially binds pre-miRNAs with a strong base-pair binding probability (BPP) in the stem region, where the mature miRNA sequence is located. Therefore, we considered a strong TRBP association to promote pre-miRNA association to Dicer (parameter *k_a_*) and therefore promote maturation efficiency (parameter *k_d_*), while antagonising dissociation (parameter *k_b_*) and stalling (parameter *k_c_*, see Figure 3B). We hypothesised that if high BPP leads to high Dicer association, and competition effects are present in pre-miRNA maturation, we would see a positive correlation between the fold change of miRNA expression following Dicer reduction and BPP (Figure 3C), as predicted by the efficient maturation of fast associating pre-miRNAs in Figure 1B and C. Conversely, if high BPP did not increase Dicer association or no competition is present in pre-miRNA maturation *in vivo* we would expect no correlation between fold change levels and BPP (Figure 3C).

We decided to use BPP for each pre-miRNA in Lee *et al.* (2011) [30] as a measure of TRBP association and therefore indirect association to Dicer. To find the BPP, the miRNA registry miRBase [28] was automatically scraped for structure information. For each miRNA, the mature sequence was extracted along with the published base pair bonds. The 5’ and 3’ strands were then aligned using RNACofold [35] and the BPP for the published bonds were extracted (see Figure 4A for the processing pipeline). The BPP for each base was then plotted against base position to provide an estimate of the stem structure (Figure 4B-C). To investigate whether BPP had any relation to miRNA expression levels, we took the area under the curve (AUC) as a single measurement and used Pearson’s correlation measure to investigate the relationship between AUC and the miRNA expression level of wild-type and YAC128 mice (Figure 4E-F).

In wild-type mice with normal *Dicer1* mRNA expression, there was a positive correlation with the *log_10_* expression level of mature miRNAs (r=0.34283, p<0.01; Figure 4E). In the YAC128 mice with significantly reduced *Dicer1* mRNA expression (Figure 4D, [30]), the correlation was also positive (r=0.43718, p < 0.001; Figure 4F). The fact that miRNA expression was positively correlated with BPP in these two independent datasets demonstrates the validity of our strategy for using BPP as a proxy measure for pre-miRNA affinity to Dicer.

To test if the relationship between the log_10_(Expression) and BPP was significantly steeper in the YAC128 than in the wild-type mice, we calculated the t-score for the regression slopes as follows:

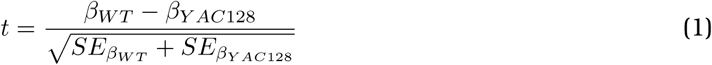

where β*_WT_* and β*_YAC128_* are the estimated regression slopes for wild-type and YAC128 mice respectively, and *SE*_β_*_WT_* and *SE*_β_*_YAC128_* are the relevant standard errors for the estimated slopes. For our model fit, the calculated slopes were 1.49±0.449 and 1.89±0.427 for wild-type and YAC128 mice respectively. These produced a t-score of t=-2.720971 with df=84, leading to a p-value of *p*=0.00791. Thus, the YAC128 mice had a significantly steeper positive association between log_10_(Expression) and BPP than the wild-type mice, is indicative of competition effects partly driving the shift in miRNA expression in the YAC128 mouse model of Huntington’s disease.

Following Dicer reduction in our model, the fold-change for all miRNAs eventually decreased (Figure 2D). When we looked at the relationship between the fold-change of YAC128 and wildtype mice we saw an overall reduction for most miRNAs, consistent with our model predictions. However, a positive correlation between fold-change and high AUC persisted (r=0.2458, p<0.05; Figure 4G). Consequently, pre-miRNAs with high BPP in the stem region are not as strongly affected by reduction in *Dicer* levels, providing further evidence for competition effects between pre-miRNA for Dicer affecting the composition of the pool of mature miRNA.

### Effects of differential pre-miRNA expression on global miRNA composition

In the previous simulations (Figure 2) we studied the effects of pre-miRNA competition for Dicer by varying Dicer concentration from low, scarce regimes to high, abundant regimes. In those simulations all eight pre-miRNA species initially had equal concentration, and differed only in their reaction kinetics. However, in real cells different pre-miRNA types likely have different abundances. This heterogeneity may have knock-on effect on Dicer competition. For example if one pre-miRNA is highly upregulated, then it may sequester more Dicer, leaving less Dicer free for other pre-miRNA types.

To investigate whether and how differential pre-miRNA expression could effect the global mature miRNA pool via Dicer competition, we returned to the same computational model used previously and successively ‘overexpressed’ (Figure 5A-B) or ‘knocked out’ (Figure 5C-D) each of our eight simulated pre-miRNAs in turn, by changing the initial pre-miRNA concentration to either 5 nM (overexpression) or 0 nM (knockout). We then ran our model with 1.55 nM available Dicer, chosen as a condition with notable competition (Figure 2C-F, G), and calculated the fold change between the mature miRNA expression in conditions with increased or knocked-down pre-miRNAs compared to the same conditions where all pre-miRNAs were expressed equally at an initial concentration of 1nM.

**Fig. 5:**
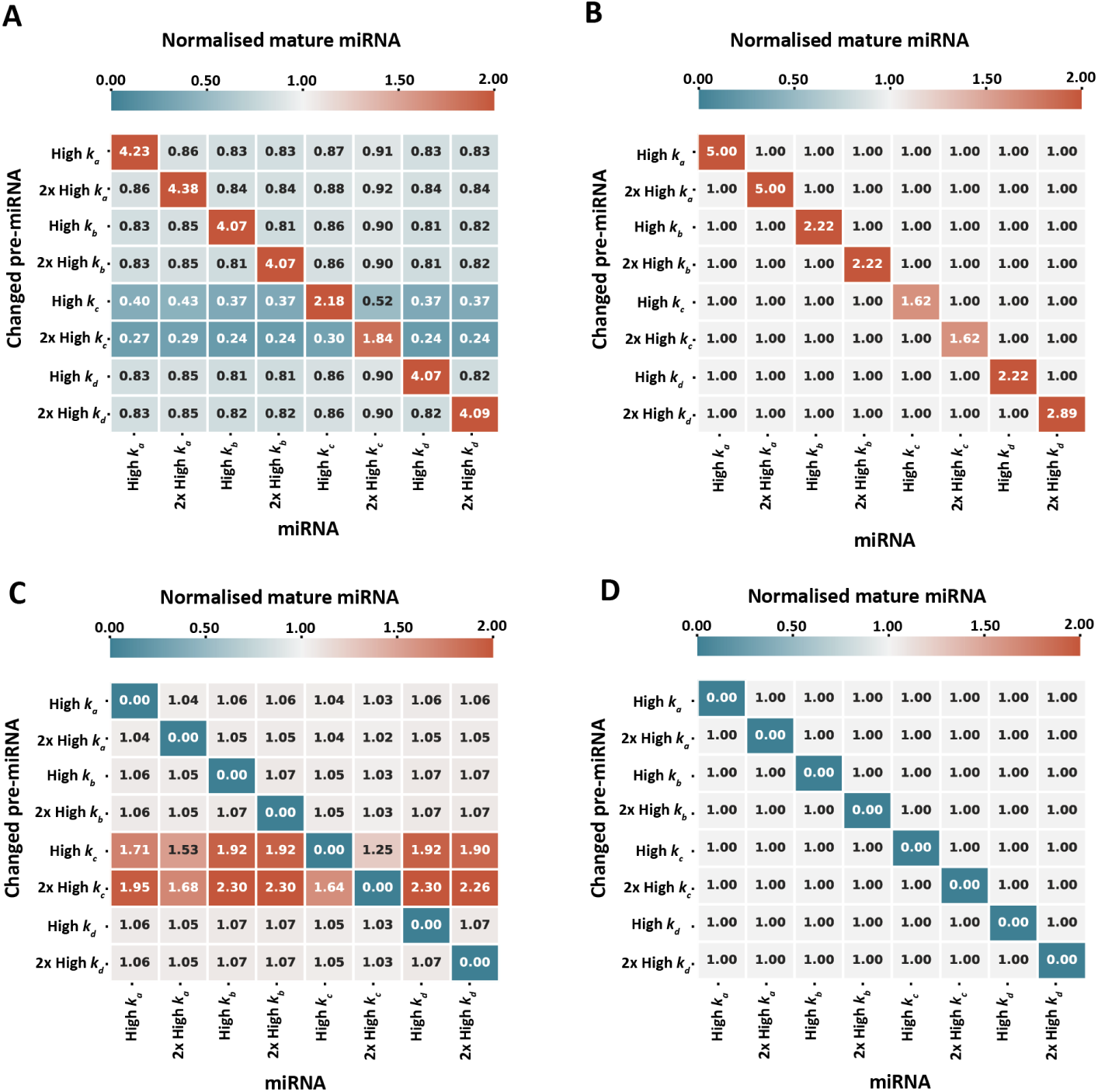
Effects of pre-miRNA increase and removal on global miRNA population. Effects on miRNA maturation in the absence (**B, D**) and presence (**A, C**) of competition following increase (5nM pre-miRNA, **A-B**) and removal (0nM, **C-D**) of specific pre-miRNAs in a system with 1.55nM available Dicer. Final concentration of mature miRNA obtained at steady state.

The simulation results are summarised in the four matrices shown in Figure 5. For a given matrix, each row corresponds to a single simulation run where a particular pre-miRNA concentration was either increased (panels A, B) or knocked out (panels C, D). The left two matrices (panels A, C) show results with 1.55 nM Dicer – encouraging competition – whereas the right two matrices (panels B, D) are the same simulations but with abundant Dicer and no competition, as a control comparison. The number and colour of each matrix element indicates the fold-change in mature miRNA expression for each of the eight species (matrix columns).

As expected, increasing the pre-miRNA concentration of a given species always increased its mature miRNA expression (red diagonals with values > 1 in Figure 5A, B). Similarly, knocking out a pre-miRNA species lead to zero mature miRNA expression (blue diagonals with values = 0 in Figure 5C, D).

Following increase of specific miRNAs, there was a broad trend of a reduction in mature miRNA of all other species in the system in a competitive environment, while the increased pre-miRNAs were all upregulated (Figure 5A). The degree of upregulation was not homogeneous between the different miRNAs, and when compared with upregulated pre-miRNAs in the absence of competition (Figure 5B) it is interesting to note that only pre-miRNAs with a fast association rate (*k_a_*) was more efficiently matured in the absence of competition when over-expressed. Increasing or decreasing the initial amount of pre-miRNAs with high *k_c_* appears to have strong effects on other pre-miRNAs in the system. While these pre-miRNAs are not highly represented among mature miRNAs (Figure 2C), their broad effects on other miRNAs in Figure 5A suggest that they play an indirect role in regulating the composition of mature miRNAs through competition for Dicer. These effects are largely mirrored following removal of each initial pre-miRNA (Figure 5C). Overall, these results demonstrate that change in expression levels of one type of pre-miRNA species can have heterogeneous knock-on effects on the expression of other miRNAs, in regimes where Dicer is scarce.

## Discussion

Modelling studies in the last few decades have discovered key principles underlying competition for resources during during gene expression and protein translation, with multiple models that investigate miRNA competition for mRNA targets [12, 23–25, 36–38] These models provide insights into the regulation of miRNA-mediated gene silencing by exploring factors that determine target specificity and affinity, allowing inferences to be made on protein expression levels. In contrast, our model does not directly concern target-specific silencing, instead we were interested in understanding the role of pre-miRNA competition for Dicer during maturation and its effect on the global miRNA population. Based on the YAC128 model of Huntington’s disease, reduction of *Dicer1* expression *in vivo* leads to a significantly altered composition of the global miRNA pool in the brain [30]. The YAC128 mouse model is also known to display a significantly altered gene expression in the brain, as determined by mRNA sequencing [39]. While the change in mRNA expression in the YAC128 mouse model might be a result of aberrant splicing activity [40, 41], tissue-specific knockdown of Dicer reveals the importance of miRNAs in regulating gene expression levels (see [42] and [43] for brain-specific examples). Additionally, Dicer expression is known to be reduced in Alzheimer’s disease [44], where changes in the miRNA transcriptome has been linked to changes in mRNA expression through complex networks [45]. Similar links between miRNA expression profiles and the mRNA transcriptome has been made in cancer ([46–48] and others) and liver failure [49]. Dicer is also reduced during ageing [44, 50], where the miRNA transcriptome is also known to be altered [51,52], highlighting the need for understanding the role of pre-miRNA competition for Dicer.

Here, we have presented a simple model of pre-miRNA competition for a limited Dicer availability during miRNA maturation. In models of competition between multiple hypothetical species of pre-miRNA, we found that high association rates provides a competitive advantage in conditions with reduced Dicer availability. We also found empirical evidence of competition effects partially regulating the composition of neuronal miRNAs in the YAC128 model of Huntington’s disease. Finally, we found that certain pre-miRNAs are more sensitive to changes in Dicer availability in conditions with increased or removed pre-miRNAs.

We showed that a strong association with Dicer (high k_a_) is beneficial both in increasing the miRNA maturation efficiency, but also provides a competitive advantage following reductions in Dicer availability (Figure 2). Based on the work of Takahashi *et al.* [33] we reasoned that pre-miRNAs with high BPP in the stem region would preferentially associate with TRBP and in turn more efficiently be loaded onto Dicer [31]. As the expression of Dicer, but not other components of the pre-miRNA maturation pathway, was significantly reduced at the mRNA level in the 12-month old YAC128 mice [30] we decided to use this as an existing model of Dicer competition *in vivo*.

We investigated the relationship between miRNA expression in the YAC128 Huntington’s mouse model [30] and BPP in the stem region. We identified a weakly positive but statistically significant positive correlation between higher expressing miRNAs and BPP in both the wildtype and YAC128 mice; though the slope of the association was significantly stronger in the YAC128 mice (Figure 4). These results provide proof-of-principle for our assumption that TRBP BPP is a valid proxy measure for Dicer affinity. We also found a weakly positive correlation in the fold change between YAC128 and wild-type mice. This shows that pre-miRNAs with a high BPP, and therefore higher TRBP association (and consequently more efficient Dicer loading and maturation), are less affected by reduction in available Dicer. These results are important because they show that competition for Dicer in part regulate the composition of the global miRNA pool. They also suggest that competition effects might play a role for the disruption of the miRNA expression profile in Huntington’s disease.

The degree of competition can not only be affected by Dicer availability, but also by differential expression of the various pre-miRNAs. We investigated this by either increasing or removing specific pre-miRNAs (mimicking up-regulation or knock-out experiments) and assessing the changes in the global mature miRNA pool in the presence or absence of competition (Figure 5). We found that pre-miRNAs with fast stalling rates were strongly affected other pre-miRNAs when over-expressed. These effects were mirrored following removal of pre-miRNAs from the system.

What might be the functional benefit of pre-miRNA competition? In evolutionary terms, MiRNAs are phylogenetically stable once they emerge. Novel miRNAs are rarely lost in descendants after arising [53]. MiRNAs are also continuously emerging and undergo changes in sequence specificity and increase sequence diversification [53]. Taken together, these suggest that competition for pre-miRNA maturation is not detrimental and could even be positive. The selective effect of pre-miRNA expression following changes in availability of pre-miRNAs with either fast dissociation or stalling rates (Figure 5) indicate that, while these inefficiently matured pre-miRNAs are not highly represented among mature miRNAs (Figure 2C, E, G), they do play an important role in shaping the composition of mature miRNAs. Competition between pre-miRNAs for Dicer might therefore help stabilise and fine-tune the mature miRNA expression.

As with all models, there are limitations with our model. First, the permanently stalled pre-miRNA Dicer complex, which can neither dissociate nor complete miRNA maturation, is not biologically plausible. While useful to account for the ceiling effect after around 40% of wild-type pre-miRNA are diced [20] (Figure 6A), there is no evidence of pre-miRNA and Dicer being removed together from the pre-miRNA maturation pathway *in vivo*. Nevertheless, our model is a single, well-mixed compartment observed for 1 hour when fitted to data. It is not unreasonable to consider the stalling a prolonged, but temporary, interruption to the pre-miRNA maturation process, for example by strong but misaligned association with a subset of pre-miRNAs within a species. In a more strongly biological model version, this term could be exchanged to *e.g.* dynamic pre-miRNA availability, spatial constraints, or including active and inactive Dicer states. Second, our model has a fixed initial concentration of pre-miRNA, and therefore does not take into account pre-miRNA production or transportation rates. However, we do not expect this to affect our conclusions, because these processes likely happen on a longer timescale of hours–days, whereas our simulations were over tens of minutes and later snapshots at steady state.

**Fig. 6:**
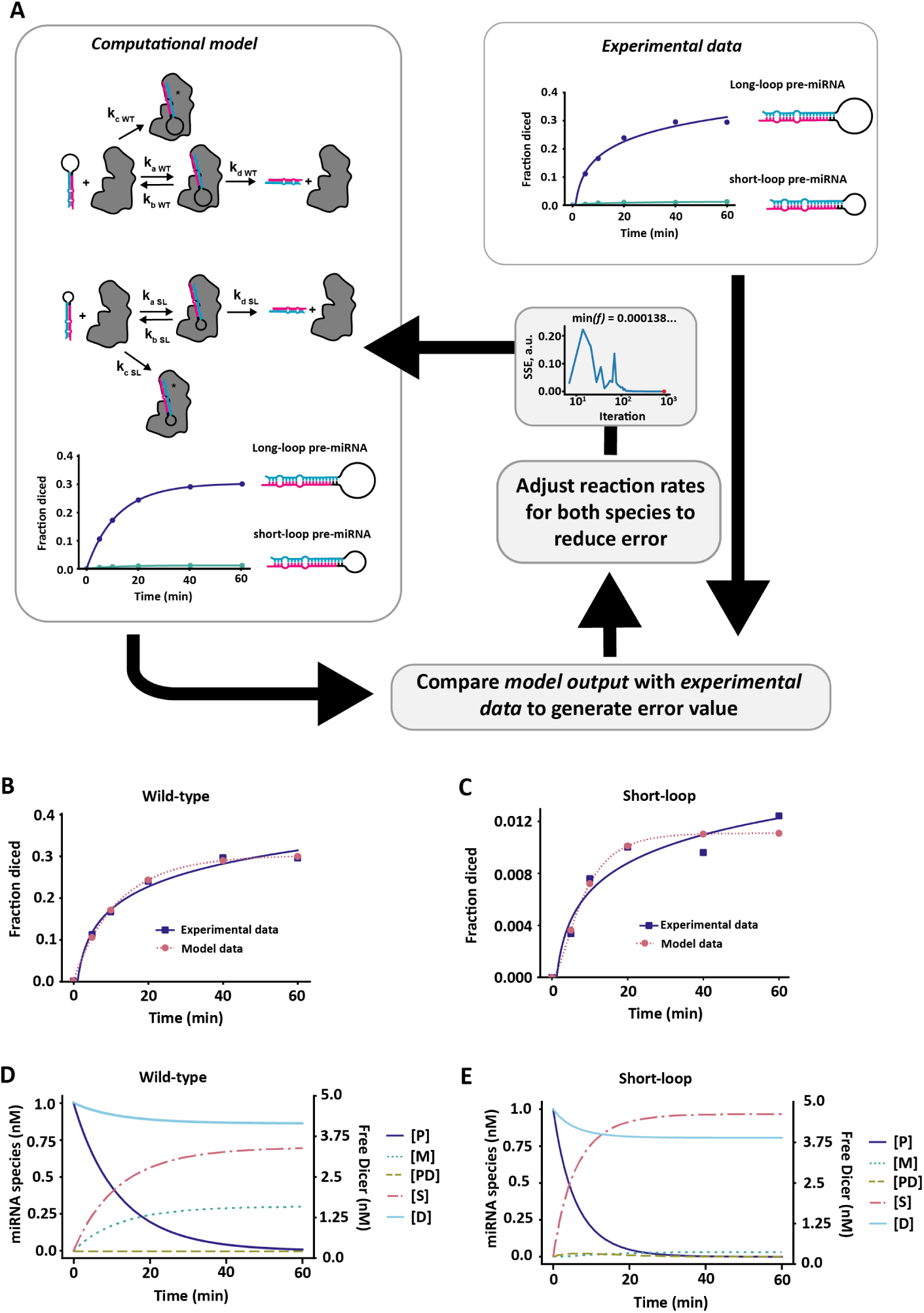
Optimisation of reaction rates in computational model. **A** Optimisation flow for fitting computational model to experimental data from [20]. The fraction of diced miRNA at specific time points in the model was compared to experimental data and the error calculated as the sum of squares (SSE) of the difference between the experimental data and data generated by the model. The error was then fed back into the optimisation algorithm CMA-ES [59] and reaction rates adjusted before output was compared again. The procedure was repeated for both wild-type (long-loop) and mutant (short-loop) pre-miRNAs until the error was maximally minimised. **B** Reproduced wild-type experimental data from model optimisation. **C** Reproduced short-loop experimental data from model optimisation. Parameter values obtained from optimisation are recorded in Table 1.

In conclusion, we have presented a simple model of pre-miRNA maturation capable of replicating experimental data. We used our model to dissect pre-miRNA characteristics that influence maturation, and based on our model predictions formed a testable hypothesis for identifying pre-miRNA competition for Dicer in experimental data. Through bioinformatic analysis we then found evidence of pre-miRNA competition *in vivo*. Finally, we uncovered a possible indirect regulatory role of lowly expressed miRNAs as directors of the global miRNA composition through pre-miRNA competition effects.

## Methods

### Software

All simulations and bioinformatics were performed in Python v3.9 if not otherwise indicated in the text. Ordinary differential equations (ODEs) were solved with the scipy integrate [54] library using the LSODA solver [55, 56], except for simulations in Figure **??**B and D where the Runge-Kutta method of order 5(4) [57, 58] was used instead to reduce numerical errors in solving. All code used in this paper is available from https://github.com/SofiaRaak/Dicer-miRNA-dynamics-model.

### Model design

We designed our initial model (Results; Figure 1B) to replicate the *in vitro* test-conditions described in Tsutsumi *et al.* (2011) [20], where 1nM pre-miRNA was incubated with 5nM recombinant *Drosophila* Dicer1 for 60 minutes and pre-miRNA maturation tracked by gel shift to determine the fraction of diced miRNA (Figure 6). Based on this, we included a set amount of pre-miRNA and Dicer in the model at the start of each simulation (Figure 6A). The pre-miRNA was then allowed to associate with Dicer to form a pre-miRNA-Dicer complex, which could either dissociate or go through a maturation step to form mature miRNA and release the Dicer again to re-join the pool. Alternatively, the pre-miRNA could irreversibly form a complex with Dicer to form a stalled, permanently sequestered pre-miRNA-Dicer species. While not necessarily biologically intuitive, this term was added to the model to account for the dynamics of pre-miRNA maturation in the experimental data.

We mathematically described our model as a series of ordinary differential equations (ODEs), where the left hand side describes the change in concentration for each species in the system and the right hand side describes the interactions and reactions that increase or decrease the concentration for each species:

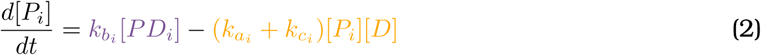

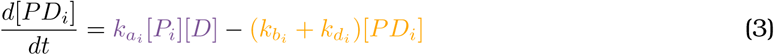

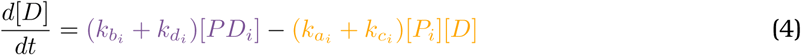

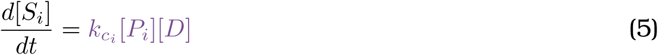

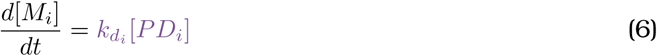

where *P_i_* represents the concentrations of free pre-miRNA of species *i*, *D* represents free Dicer, *PD_i_* represents pre-miRNA bound to Dicer, *S_i_* represents stalled pre-miRNA-Dicer complex and *M_i_* represents mature miRNA concentration (all in nM). The reaction rates *k_a__i-di_* represent the reaction rates for pre-miRNA-Dicer association, dissociation, stalling and dicing respectively, see table 1. The model was then fitted to experimental data.

Although dissociation constants are often more interpretable than reaction rates, in multi-reaction systems, especially those with sinks like the model we present, knowledge of the dissociation constant for the reactants in any given bimolecular reaction does not necessarily imply knowledge of the resulting steady-state concentration of the product. As a consequence, in this study we chose to describe the model in terms of its more elementary reaction rates.

### Fitting model to data

In order to fit the parameters of the model to experimental data initial parameter values were specified and the system of ODEs solved and output compared to the experimental data in Tsutsumi et al [20]. We constrained the ratio of the *k_a_* and *k_b_* reaction rates to match Tsutsumi et al’s [20] reported *K_d_* values for both WT (*K_d_*= 25.4 nM) and short loop mutants (*K_d_*= 147.7 nM) but fit all other parameters. The difference between the model output and experimental data was then quantified as the sum squared error (SSE) between the simulation time series and the experimental data time series and fed into the CMA-ES [59] optimiser, iteratively updating the parameter values and repeating the process until a good fit was achieved (see fig 6). The optimised parameter values for both wild-type (WT) and mutant (short-loop; SL) miRNA are shown in table 1, along with initial concentrations of pre-miRNA (P_0_) and Dicer (D_0_). In all subsequent simulations, the WT parameters were used as a foundation for the different pre-miRNA species.

**Tab. 1:**
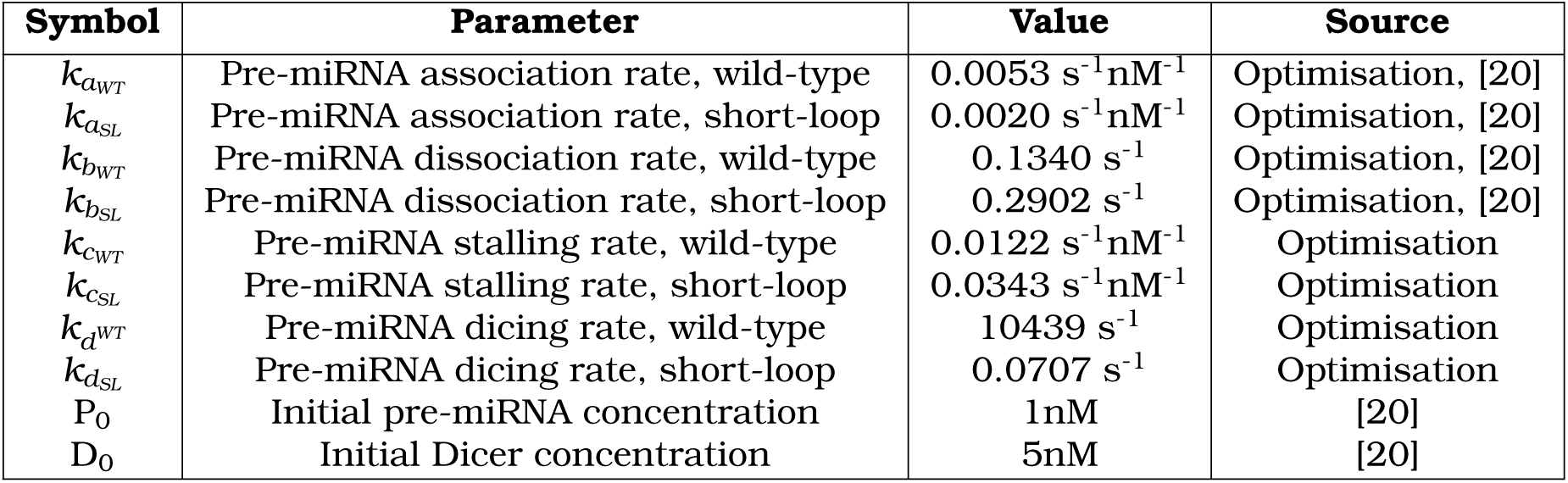
Parameter values used in model.

### Bioinformatics

All miRNA expression data used in bioinformatics are available in [30]. Briefly, miRNAs present in expression data from wild-type and YAC128 mice were identified in miRBase [28] and their pre-miRNA structure recorded. MiRNAs that were not found in miRBase or that did not have a pre-miRNA structure were excluded. From the pre-miRNA structure, the mature sequences were isolated and their base pair bonds recorded. The mature sequences were then aligned with RNACofold v. 2.4.13 [35] and the base pair binding probability (BPP) for the recorded base pair bonds were extracted. In the absence of a recorded base pair bond, the mean BPP was chosen (Results; Figure 4A). The BPP was then plotted against the nucleotide position to create a BPP curve (Results; Figure 4B). As a single measure of BPP, the area under the curve was taken and plotted either against the log_10_(miRNA Expression) or the fold-change in miRNA expression between wild-type and YAC128 mice (Results; Figure 4).

